# Online supervised learning of temporal patterns in biological neural networks under feedback control

**DOI:** 10.1101/2025.08.10.669034

**Authors:** Yuki Sono, Hideaki Yamamoto, Yusei Nishi, Takuma Sumi, Yuya Sato, Ayumi Hirano-Iwata, Yuichi Katori, Shigeo Sato

## Abstract

In vitro biological neural networks (BNNs) provide a well-defined model system to constructively investigate how living cells interact with their environment to shape high-dimensional dynamics that could be used to generate a coherent temporal output, such as those required for motor control. Here, we developed a real-time closed-loop BNN system capable of generating periodic and chaotic temporal signals by integrating cultured cortical neurons with microfluidic devices and high-density microelectrode arrays. We show that training a simple linear decoder with fixed feedback weights enables the system to learn and autonomously generate diverse temporal patterns. When feedback was switched on, irregular activity in BNNs is transformed into low-dimensional, structured dynamics, producing coherent trajectories characterized by stable transitions between neural states. BNNs trained on different target frequencies—ranging from 4 to 30 s—could be trained to sustain oscillations at distinct frequencies, demonstrating their adaptability. Importantly, a top-down control of self-organized network formation with microfluidic devices is the key to suppress excessive synchronization and increase dynamical complexity in BNNs, facilitating the training and robust output generation. This work offers a biologically inspired platform for understanding the physical basis of cortical computation and for advancing energy-efficient neuromorphic computation.

**Significance Statement:** Reservoir computing is a machine learning paradigm that exploits the transient dynamics of high-dimensional nonlinear systems. Although it was originally inspired by the mammalian brain and widely explored in physical systems, its implementations in biological neural networks (BNNs) have been limited due to their excessive connectivity and global synchrony in vitro. Here, we use microfluidic devices to construct modular, nonrandomly connected BNNs and integrate them with microelectrode arrays in a closed-loop reservoir computing environment. We show that the system can be trained to autonomously output various temporal signals, with the modular connectivity that is essential for learning. In vitro BNNs provide unique alternatives for physical reservoirs with dynamic adaptability.

## Introduction

Precise motor action control is essential for animal survival. For this purpose, an animal uses the intrinsic chaotic neural activity that arises from the recurrent neural network (RNN) in its brain as a substrate to generate coherent muscle commands^1,2^. Importantly, the brain does not operate as a simple open-loop controller; instead, muscles continuously provide sensory feedback to the brain, which stabilizes motor actions and enables adaptive behaviors in dynamic environments^3^. Computational studies have previously demonstrated that such computations can be modeled with RNNs by incorporating feedback from the output layer and that stable learning can emerge by tuning these output-feedback systems^4–6^. Nevertheless, the biological plausibility of these mechanisms and whether such principles can be realized in biological neural networks (BNNs) consisting of living neurons remain largely unexplored.

Recently, in vitro BNNs integrated with reservoir computing models^7,8^have emerged as unique platforms for constructively investigating how living neural systems can perform complex computations^9–12^. This approach is also compelling in the context of physical reservoir computing, which is a form of neuromorphic computing that harnesses the transient, high-dimensional dynamics of nonlinear physical systems for computation^13–15^, since individual neurons (as computational units) exhibit complex, nonlinear input–output transformations^16^. Furthermore, BNNs distinguish themselves from other physical reservoirs in several key ways. For example, unlike conventional reservoir computing systems that require stable internal dynamics, BNNs can exhibit plastic changes in their responses over time^17–20^. Counterintuitively, this property can be leveraged to enhance the classification and prediction performance of temporal signals^12^. Beyond the applications of BNNs in reservoir computing, their plasticity may enable them to be tuned under the framework of free-energy principles^21^ or reinforcement learning^22^ to optimally interact with the environment.

Another hallmark of BNNs is their ability to exhibit spontaneous activity, which is persistent background activity that occurs even without external inputs^23–25^. In the mammalian cortex, such spontaneous activity plays critical roles in the network formation^26^, memory consolidation^27^, and prediction^28^. This activity is also preserved even when cortical neurons are cultured in vitro^29^. Theoretical studies have shown that feedback signals can stabilize chaotic dynamics in neural networks, enabling the generation of coherent time series signals from networks exhibiting irregular spontaneous activity^4,30^. However, testing the biological plausibility of such mechanisms in the context of BNN-based physical reservoir computing requires the integration of several demanding technologies, such as the preparation of in vitro BNNs with high-dimensional dynamics and the development of closed-loop controllers with sufficient spatiotemporal resolutions.

Here, we integrated microfluidics-based cell patterning with CMOS-based high-density microelectrode array (HD-MEA) recording to realize closed-loop reservoir computing with BNNs that autonomously generates coherent temporal signals. Microfluidic devices were used to constrain the growth of cultured rat cortical neurons in a nonrandom architecture resembling the modular connectivity in the mammalian cortex and to suppress excessive synchronization across the network. The HD-MEA was then used to record the neuronal activity at a high spatiotemporal resolution, which was streamed to a desktop computer for real-time processing and used to generate feedback stimuli for the BNN. We show that micropatterned BNNs can be trained online to generate a variety of periodic and chaotic time series signals. Importantly, this was possible only when BNNs were patterned to bear modular connectivity, which prevented the formation of excessively dense connections. Our results offer a framework for understanding how animals may transform chaotic neural dynamics into coherent motor commands while also highlighting the potential of BNNs as novel computational resources.

## Results

### Closed-Loop Control Using HD-MEAs

A closed-loop controller for BNNs was constructed using an HD-MEA system (Fig. 1A). The system recorded spike trains from the electrodes, filtered the spike trains with a double exponential kernel to convert them into a continuous signal (a reservoir state **x**(*t*)), decoded the output signals via a linear readout defined by **y**(*t*) = **W**(*t−* Δ*t*)**x**(*t*), and sent a feedback stimulation to the BNN. Here, **W**(*t−*Δ*t*) is the output matrix at a previous time step, which was trained via first-order reduced and controlled error (FORCE) learning^4^ to minimize the error between **y**_*t*_ and the target signal. The feedback stimulation with an amplitude of **A**(*t*) = *g*(**y**(*t*)) was delivered to selected electrodes, where *g* is a mapping function (described in the Methods section).

**Fig. 1:**
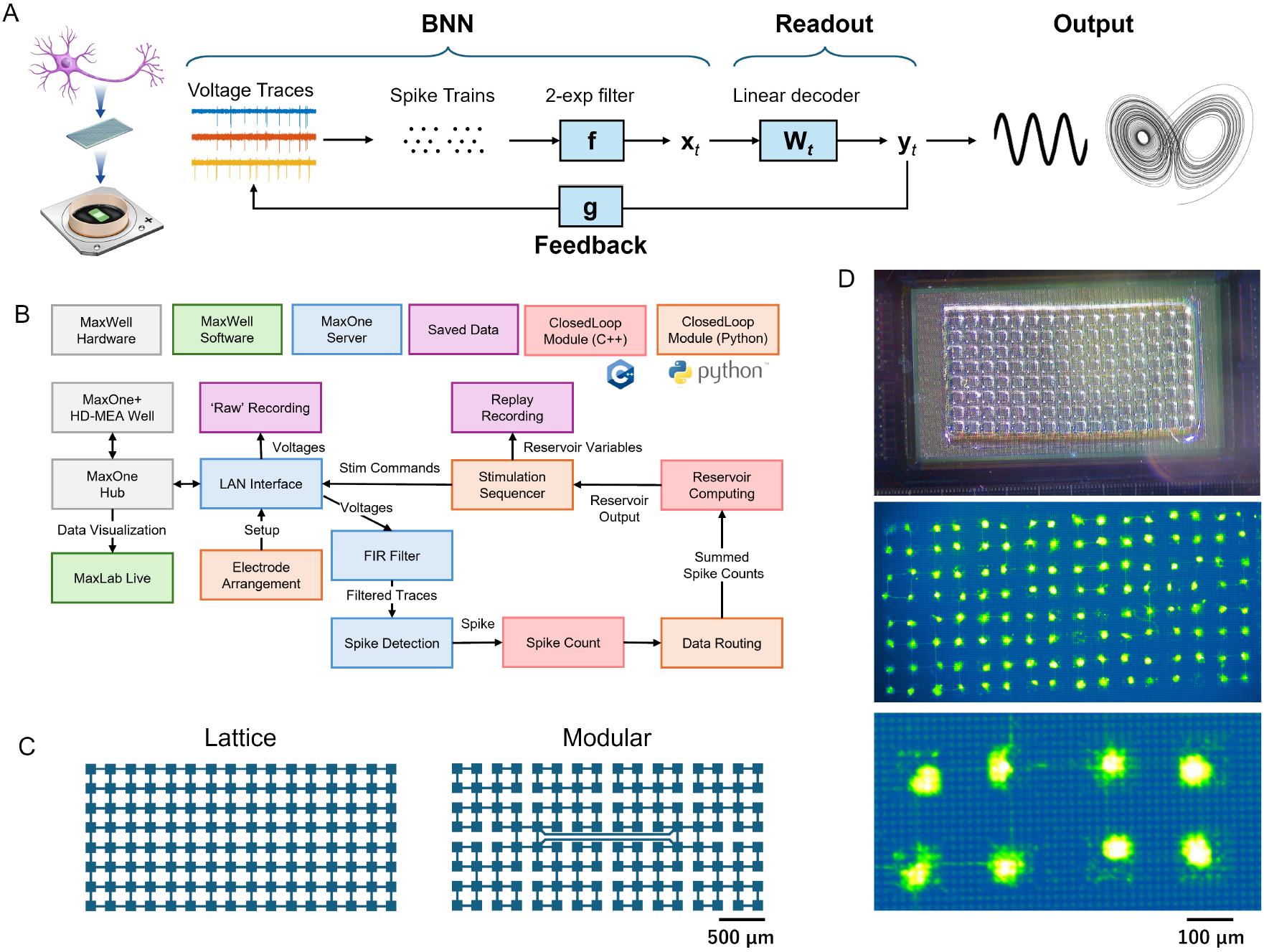
Overview of the experimental setup. (A) Signal generation task. Voltage traces recorded from the BNN were converted into spike trains, which were then passed through a double exponential filter to generate a continuous-time signal, referred to as the reservoir state. The readout layer transformed the reservoir state into an output signal, which was then used to generate feedback input for the BNN. (B) Data flow in the closed-loop setup. The detected spikes were passed on from the hardware to the Python application programming interface (API), and they were then used to perform the reservoir computing task and compute the feedback stimulation in a custom Python/C++ script. The obtained stimulation signal was then transferred to the hardware via the API. (C) Structures of the lattice and modular networks. Both patterns contained the same number of wells in which neuronal cell bodies attached but differed in the designs of their interconnecting microchannels, which shaped the network topology. (D) Recording area of the HD-MEA with the PDMS-microfluidic device attached (top). Primary cortical neurons at DIV 29 bearing a modular pattern (stained with the neuronal marker NeuO) are shown in the middle and bottom panels.

Technically, the entire system was realized by controlling the HD-MEA in real time with custom-written C++/Python scripts (Fig. 1B). The C++ script received instantaneous spike counts from the hardware, which sent summed spike counts for each electrode to a Python closed-loop controller. The Python controller then routed the data to a second C++ script for filtering, decoding, and output training. The decoded output was then passed to the Python controller, which created a stimulation sequence that was subsequently sent back to the BNN through the HD-MEA. The full implementation details are provided in the Materials and Methods section.

The BNN was prepared by culturing primary rat cortical neurons on HD-MEA chips. The HD-MEA contained 26,400 electrodes aligned in a 17.5-µm pitch over a 2.1×3.9-mm^2^ recording area. Unlike sparsely coupled networks that are used as reservoir layers in reservoir computing models, cultured neurons tend to form dense interconnections when grown on a homogeneous substrate^31,32^. To restrict excessive neuronal connections, a homemade polydimethylsiloxane (PDMS) microfluidic film was placed on the electrodes to physically constrain the adhesion and growth of the neurons and their neurites^20,33,34^. Two designs were fabricated: *lattice* and *modular* versions (Fig. 1C). Both designs contained 128 square wells with areas of 100*×*100-µm^2^, which were arrayed across the entire recording area of the HD-MEA. Each well contained a population of neurons that formed aggregates and was connected with the microchannels penetrated by the neurites to functionally couple the neurons in wells (Fig. 1D). The two configurations differed in their interwell connectivity, with the modular design featuring sparser connections than those of the lattice design. BNNs grown without the microfluidic film, denoted as *homogeneous*, were used as a control.

### Spontaneous Activity of BNNs

First, the intrinsic properties of the BNNs were assessed by recording the spontaneous activities derived from homogeneous, lattice, and modular BNNs (Fig. 2A). The homogeneous networks exhibited bursting patterns with high degrees of synchronization across the entire chip, suggesting that the neurons formed a highly interconnected network without a distinct spatial organization. In contrast, the spontaneous activity of the lattice and modular networks was characterized by spatially segregated bursting patterns, reflecting the constrained connectivity imposed by the microfluidic devices.

**Fig. 2:**
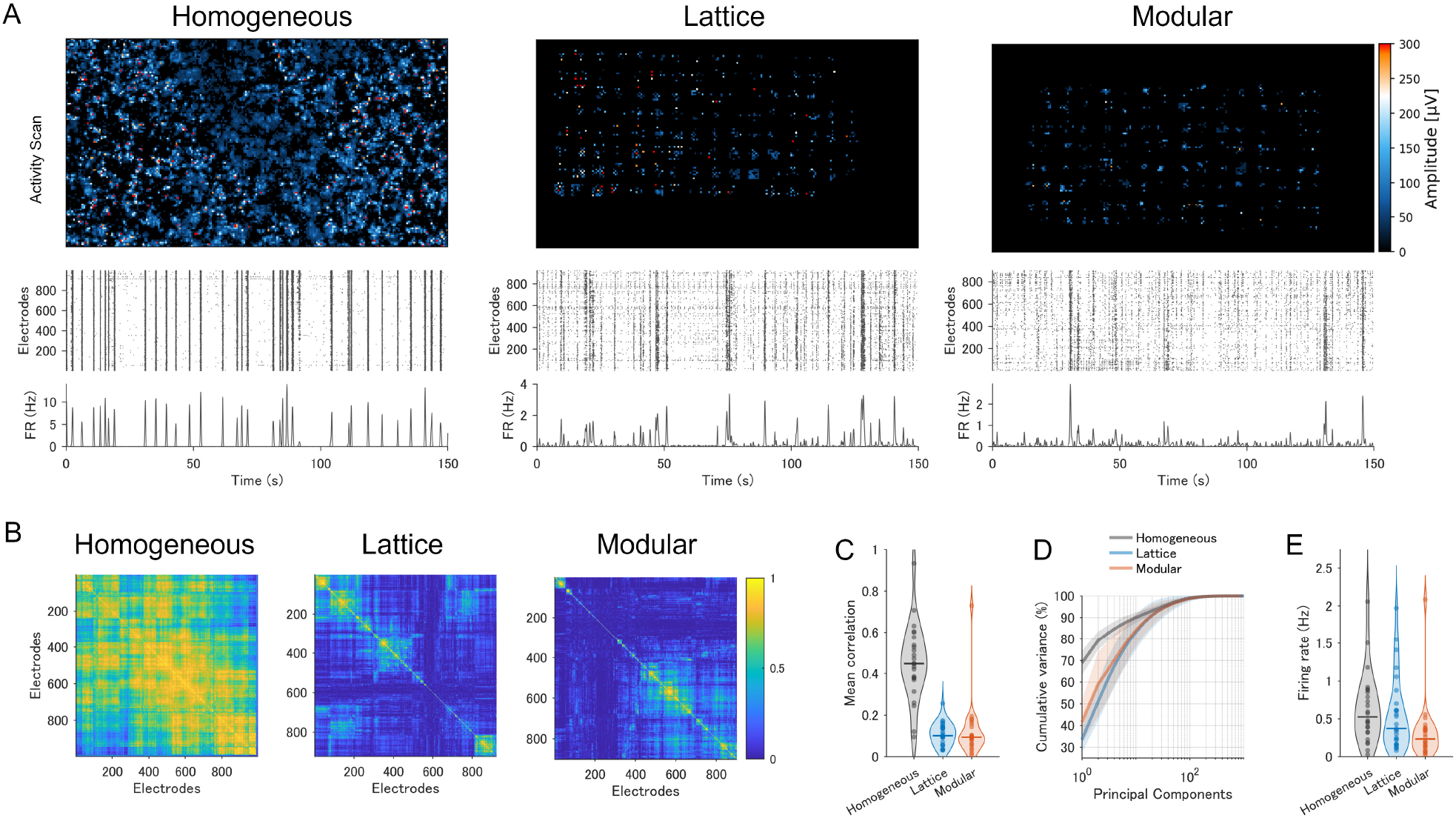
Spontaneous activity of homogeneous and engineered BNNs. (A) Live-cell electrical images of each network, which were generated by scanning across the electrode array and color-mapping the mean amplitude of the detected presumptive action potentials (top). For the lattice and modular networks, only the electrodes within the well regions were scanned. Representative raster plots (middle) and corresponding population firing rates (bottom) are also shown for each network topology. The data were obtained from cultures at DIV 14. (B) Pairwise correlation matrices computed from the same networks shown in (A). (C) Network-averaged mean correlations for the three topologies. Each plot represents a single BNN realization, and the horizontal bars are their medians. (D) Distribution of the variance explained by each principal component. The shaded error bars represent the standard deviations. (E) Comparison among the mean firing rates for the three topologies. Each plot represents a single BNN realization, and the horizontal bars are their medians. Sample sizes: *n* = 24 (homogeneous), *n* = 27 (lattice), and *n* = 24 (modular).

An analysis of the network statistics revealed that the mean pairwise correlations were significantly lower in both the lattice and modular networks than in the homogeneous culture (mean*±*sem): 0.45*±*0.04 (homogeneous, *n* = 24), 0.11*±*0.01 (lattice, *n* = 27), and 0.12*±*0.03 (modular, *n* = 24) (Figs. 2B, C). This reduction in synchrony increased the dimensionality of the network dynamics, which were characterized by broader variance distributions across the principal components^35^ (Fig. 2D).

The most prominent difference between the lattice and modular networks was a nearly twofold increase in the firing rate in the lattice configuration relative to the modular configuration (*p <* 0.05, one-sided *t* test), reflecting the influence of abundant intermodular connections: 0.63*±*0.10 Hz (homogeneous, *n* = 24), 0.56*±*0.09 Hz (lattice, *n* = 27), and 0.32*±*0.08 Hz (modular,*n* = 24) (Fig. 2E). Throughout the development process (14–31 days in vitro (DIV)), there was a weak trend for the mean firing rate to increase and the mean correlation to decrease, yet the overall effect of microfluidic patterning remained stable (Supplementary Fig. S1). Taken together, these observations underscore the ability of engineered BNNs, especially lattice networks, to support flexible dynamics with high activity levels, providing a preliminary indication of their potential to function as effective reservoirs.

### Periodic Wave Generation Task

Next, as a basic example of online training in a BNN-based reservoir computing task, the neural activity was modulated by the feedback controller (see Fig. 1A), whose readout weights were trained in real time via the FORCE algorithm. Sinusoidal waves with periods of 4, 10, and 30 s were used as the target waves, and the performances of lattice, modular, and homogeneous BNNs were compared.

Fig. 3A illustrates how the BNN activity can be modified so that it autonomously produces a periodic, sinusoidal output. Initially, without any external drive, the neurons in the BNN exhibited chaotic spontaneous activity, as detailed in the previous section. When a feedback input was added and FORCE learning started, the readout weights began to fluctuate rapidly, which immediately caused the neurons in the BNN to activate periodically and forced the output to match the target sine wave. Over time, the fluctuations exhibited by the readout weights decreased, and a quasistatic weight matrix that generated the target function was obtained (Fig. S2). At this point, the learning could be turned off, and then the BNN with the feedback drive sustained the oscillatory activity and the output. The learned oscillatory dynamics were not always robustly retained in the postlearning phase, and the mean squared error (MSE) increased in the postlearning phase relative to the learning phase in 99% of the trials (*n* = 143; all network topologies pooled; target signal: sine wave with *T* = 30 s). Nonetheless, the same approach could be used to apply BNNs to generate other periodic signals that contain multiple frequency components and nondifferentiable points, such as triangle wave or square waves (Fig. 3B). Hence, FORCE learning, which has been shown to be successful for training rate-based and spiking neural networks (SNNs)^4,36^, can be used to train BNNs to generate various temporal signals.

**Fig. 3:**
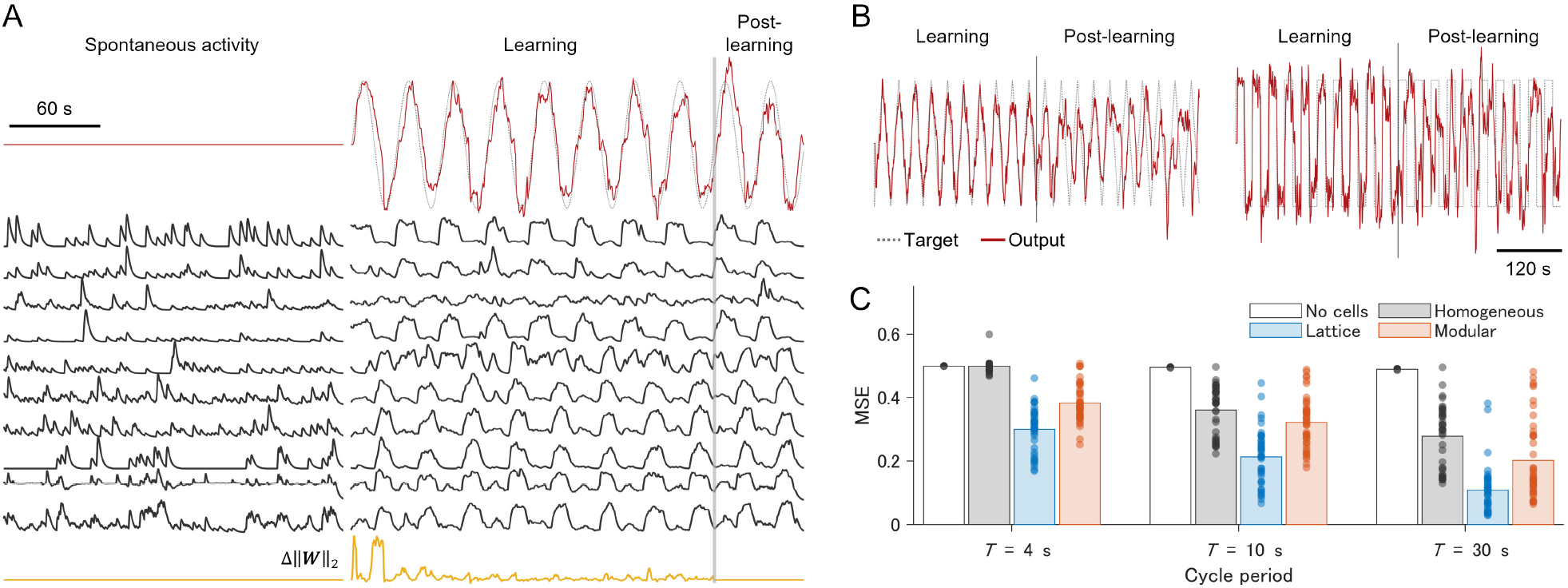
FORCE learning in BNNs. (A) FORCE training sequence in a modular network at DIV 23, presented as an homage to Figs. 2A–C of Sussillo and Abbott^4^. The network output is shown in red, whereas the target signal, a sinusoidal wave with a cycle period of 30 s, is shown in dotted gray. The black traces are the reservoir states (filtered spike trains) derived from the ten electrodes with the largest readout weight magnitudes. The yellow trace is the time derivative of the magnitude of the readout weight vector. After learning, the BNN was able to sustain the trained oscillation even after the weight update was disabled. (B) Examples of FORCE learning applied to other periodic waveforms: a triangle wave (left) and a square wave (right), both with cycle periods of 30 s. (C) Comparison among the MSEs induced during the learning phase across different sine-wave periods and network configurations. The bars represent the means, and single BNN simulations are plotted.

The generation of such temporal signals requires a BNN to exhibit complex, asynchronous responses to feedback stimulation (Fig. 3C). For a target signal with faster oscillations (a sine wave with a 4 s cycle), the MSE during the learning phase of the homogeneous networks was indistinguishable from that of a random output that was generated with no neurons (two-sided *t* test, *p* = 0.53). The MSEs of the homogeneous networks decreased when the period of the target signal was increased but were consistently higher than the values obtained from the lattice and modular networks. Comparing the two types of micropatterned networks, the lattice networks consistently presented lower MSEs than the modular networks did, presumably reflecting their greater degree of network activity, facilitating regression by the linear decoder. The finding that the regression performance improves for target sine waves with longer cycle periods is consistent with a previous report showing that a homogeneous BNN can be trained via FORCE learning to generate a constant, time-invariant output^10^.

### Network Dynamics and Weight Matrices

To better understand the mechanism behind the increased regression performance achieved by the engineered BNNs, we analyzed the network dynamics during the learning phase. As in the case with spontaneous activity (see Fig. 2A), the homogeneous networks exhibited irregular bursting activity that spread across the entire networks even under closed-loop control. This led to consistent failures of the FORCE training for all target sine waves in the homogeneous networks (Fig. S3A). Although the decoded output *y*(*t*) fluctuated in response to the network activity, these fluctuations occurred at random time points, and the output lacked the smooth, periodic profile of a sine wave.

This behavior contrasted with that observed in engineered networks (Figs. S3B–C). For example, in the lattice networks, the reservoir state was modulated by a feedback controller to exhibit periodic changes in sync with the target signal. These dynamics enabled the regression of output signals that closely tracked the target waveform, demonstrating the ability of BNNs to be used for generating coherent time series signals. Importantly, the same culture preparation could be trained to sustain oscillations with arbitrary cycle periods after terminating the learning process (Figs. 4A–C).

**Fig. 4:**
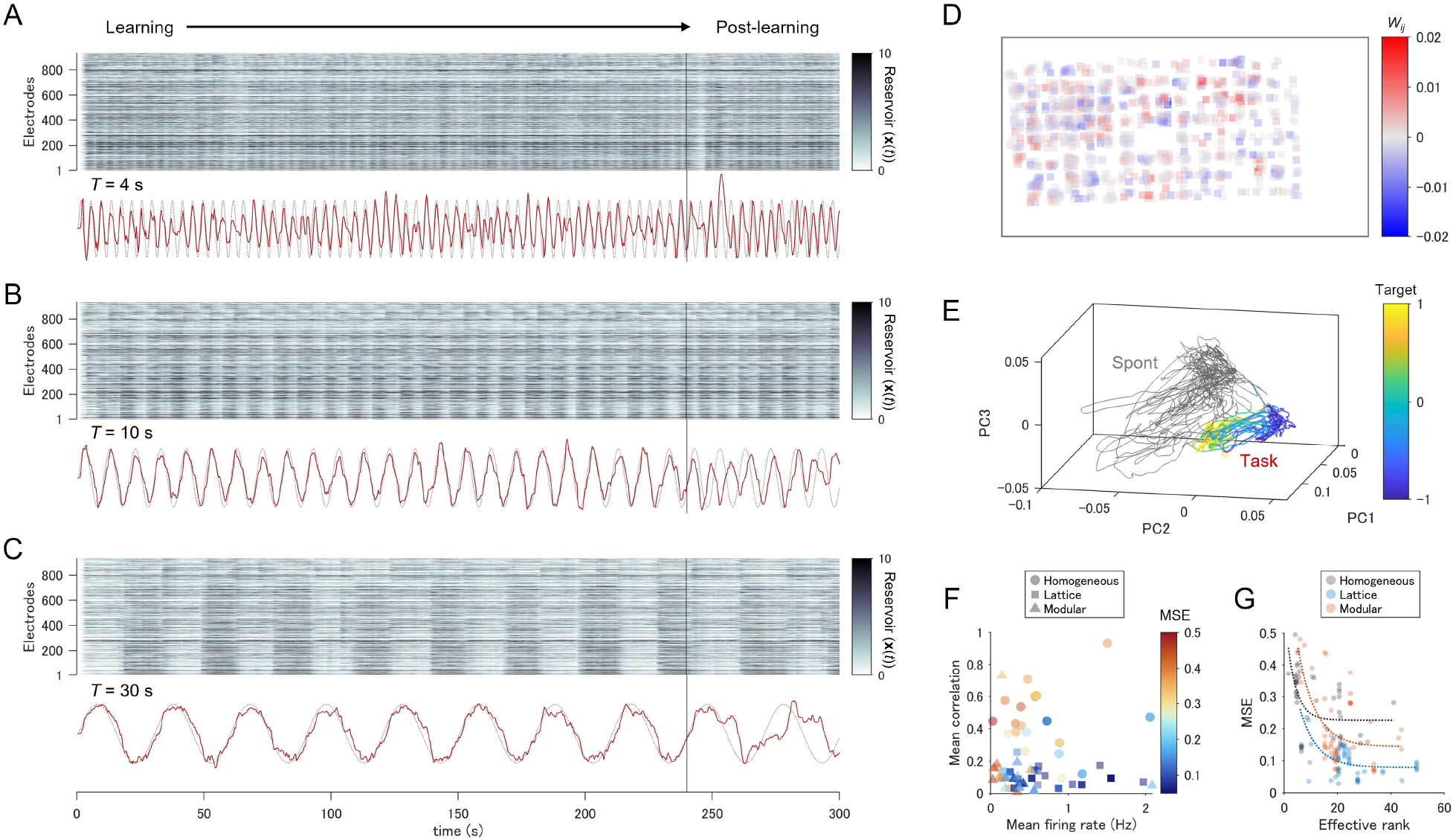
Versatility of BNN reservoirs and their supporting network dynamics. (A–C) Reservoir states derived from all electrodes (top) and the corresponding output signals (bottom), obtained from an identical lattice network at DIV 24. The cycle periods of the target sinusoidal wave were set to 4 s (A), 10 s (B), and 30 s (C). Note the difference in the frequency of the output oscillations after the weight update was halted, reflecting the ability of the system to autonomously sustain temporal dynamics across different timescales. (D) Spatial distribution of the learned output weight vector from the same network shown in (A–C). (E) Trajectory of the reservoir states projected in the principal component space. The gray trace represents the spontaneous activity prior to learning, whereas the colored trace represents the trajectory during the process of learning a sinusoidal wave with a 30-s period. The trace color encodes the amplitude of the computed reservoir output, illustrating the correspondence between the neural state space and the output signal. (F, G) Effect of network dynamics on the MSE during the learning phase for a 30-s sinusoidal wave. The mean firing rate, mean correlation, and effective rank were evaluated during spontaneous activity. Each plot represents a single recording. Sample sizes: *n* = 42 recordings from 7 cultures at DIV 14–28 (homogeneous), *n* = 52 recordings from 7 cultures at DIV 14–30 (lattice), and *n* = 49 recordings from 8 cultures at DIV 14–30 (modular).

An analysis of the readout weight distributions further revealed that the output signal **y**_*t*_ was reconstructed through contributions from the entire network, which was consistent with the spatially complex dynamics observed in the engineered BNNs (Fig. 4D). Moreover, a trajectory analysis revealed that when feedback control was activated, the network dynamics rapidly transitioned from a chaotic state (gray) to a more structured state (colored), in which the system transitioned between two attractors that corresponded to the positive and negative peaks of the sine wave (Fig. 4D).

Fig. 4F summarizes how the regression performance depends on the firing rate and neural correlations within the BNN. Regardless of their network morphologies, BNNs with higher mean firing rates in their spontaneous activity exhibited lower MSEs during the learning phase. A high mean correlation was associated with poor performance. However, a low mean correlation did not necessarily guarantee high performance, as the MSEs in this range were highly variable. Overall, the best regression performance was thus observed in BNNs with high firing rates and low correlations.

A clearer inverse relationship was observed between the effective rank of the network activity and the MSE (Fig. 4G). The effective rank increased when the eigenvalues obtained via principal component analysis (PCA) were distributed more broadly into higher PCs (see Fig. 2D)^37^. Since a broader eigenvalue distribution implies that the corresponding network exhibits more complex and higher-dimensional dynamics^35^, the result suggests that networks with richer internal dynamics tend to support a more accurate regression.

Collectively, these results underscore the importance of the dynamical complexity enabled by microfluidic patterning in the use of BNNs for functional computation.

### Chaotic Wave Generation Task

Finally, we tested the applicability of the FORCE-trained BNN to learn to generate a Lorenz attractor, which is a nonlinear chaotic time series (Fig. 5A). The input layer was extended to handle three dimensions, corresponding to the *x, y*, and *z* components, and the values of each component were encoded as input pulses via pulse amplitude modulation^38^. A simple linear decoder, whose weights were trained by the FORCE algorithm, was used for decoding.

**Fig. 5:**
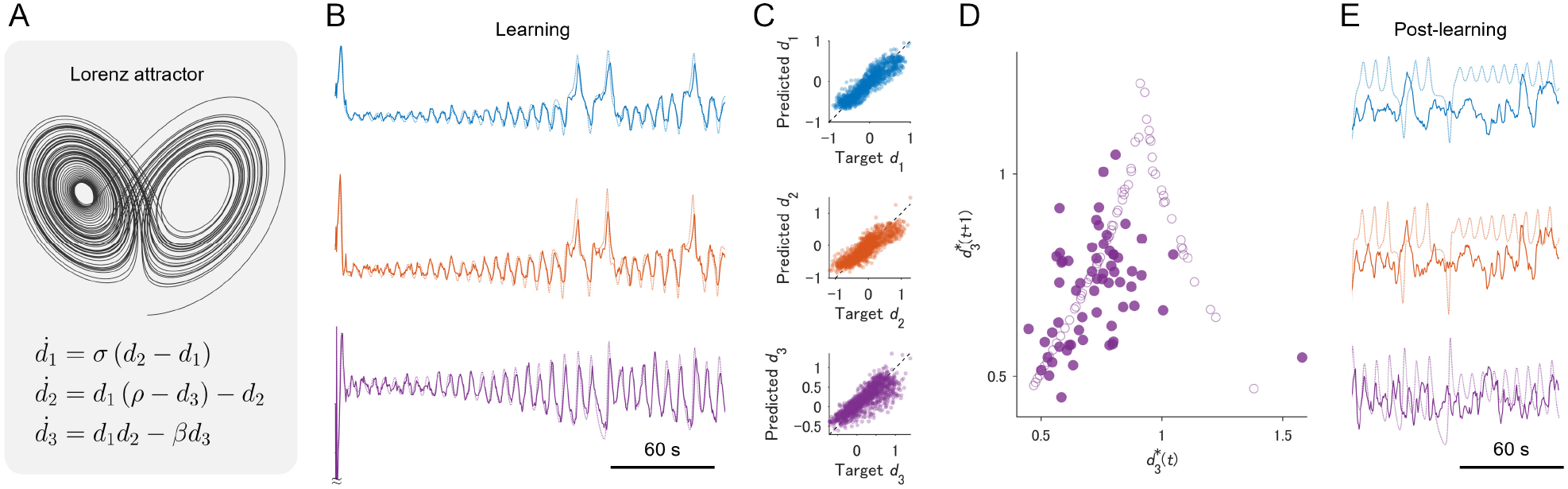
Learning a nonlinear chaotic equation. (A) The trajectory and governing equation of the Lorenz attractor. The actual target signal was obtained by reshaping the waveform, as detailed in the Methods section. (B) Time series of the *x, y*, and *z* components during learning, using a lattice network at DIV 24 as the reservoir layer. (C) Point-wise comparison between the target and predicted signals over time; each dot represents a sample point in time. (D) Lorenz map generated from the target signal (open circles) and the actual network output (filled circles). The axes indicate the peak amplitudes for the *z*-component at the *t*th and (*t* + 1)th time steps. (E) Output of the same network as that shown in (B) after the FORCE learning was halted.

Similar to the periodic wave generation task, the engineered BNN, but not the homogeneous BNN, could be successfully trained to follow the target trajectory during the learning phase (Fig. 5B). When comparing the predicted and target signals over time, their pairwise correlations exceeded 0.8 in all dimensions, indicating an accurate fit (Fig. 5C). A further analysis of the generated trajectory revealed that the system was especially successful at reproducing signals in the low-amplitude range (Fig. 5D). The reduced accuracy observed in the high-amplitude range was likely due to the current setting of the *α* parameter in the FORCE learning; although lowering this parameter could yield improved performance, it would increase the risk of signal divergence. Finally, after the online weight update was halted, the output trajectory began to deviate from the target signal (Fig. 5E). Nonetheless, the system preserved its intrinsic dynamical structure, continuing to generate a chaotic oscillatory output. These results demonstrate that engineered BNNs can be trained to generate complex nonlinear dynamics under closed-loop feedback control, even under biologically constrained conditions.

## Discussion

This study demonstrates that BNNs can be trained in real time via FORCE learning to autonomously generate structured temporal patterns, bridging in vitro neurodynamics with machine learning. While biological neurons inherently possess the capacity to self-organize and form functional networks even in vitro, the resulting networks tend to exhibit excessively synchronized bursting behaviors. Bioengineering technologies involving microfluidic devices offer an effective approach for bridging the gap between in vitro and in vivo BNNs^20,39^. In this context, we demonstrated that rat cortical neurons cultured in lattice and modular configurations developed into networks that exhibited more diverse and structured activity patterns that could be used as the reservoir layer for learning various periodic waveforms, as well as more complex chaotic dynamics.

FORCE training was initially introduced as a framework to extract diverse temporal patterns from artificial neural networks composed of rate neurons^4,6^and was later shown to be applicable to SNNs^36,38^. Its application to BNNs was first demonstrated by Yada et al.^10^, who reported that the spontaneous activity in homogeneous BNNs can be controlled using FORCE learning to produce a constant output signal via a linear decoder. Our work extends this framework to enable the generation of temporally varying signals, which was accomplished by enriching the internal dynamics and stimulus responses of the in vitro BNNs by micropatterning.

One current limitation of this work is the decreased performance observed during the postlearning phase, when the weight updates were halted and the system ran autonomously. This could have arisen from the nature of FORCE learning, which tends to be fragile to reservoir perturbations. For example, in a previous involving using SNNs, removing as few as 10% of the reservoir neurons resulted in a catastrophic decrease in task performance^36^. In theory, the *α* parameter in Eq. 11 (see Methods section) scales the regularization term in the FORCE learning, which is based on recursive least squares^40^; thus, increasing the value of this parameter would suppress overfitting. However, increasing *α* also slows the learning procedure and hampers the generation of high-amplitude signals. Consequently, its value was heuristically set to 1,000 in the present work to balance these trade-offs. Recent extensions of FORCE learning, including transfer-FORCE and composite-FORCE, may also improve the performance of BNN-based reservoir computing by increasing the convergence speed to compensate for the limited timescales of biological preparations and by stabilizing the weight updates under noisy conditions^41,42^. Furthermore, in a separate study using SNN simulations, we recently reported that the feedback delay critically impacts the performance of a time series generation task^38^. Since a large fraction of the feedback delay in the preset system originates in the signal filtering step prior to performing spike detection, the delay could be reduced by using specialized hardware^43^ or changing the filtering algorithm^44^, which could further improve the overall task performance and increase the computational capacity of the closed-loop controlled BNN reservoir.

A promising future direction is related to the application of brain–machine interfaces and neuroprosthetic devices^45^. In particular, the ability to modulate and decode neural dynamics in real time via closed-loop interactions may offer new strategies for augmenting or restoring motor and cognitive functions. Moreover, in vitro cell cultures, especially those derived from human-induced pluripotent stem cells, are increasingly gaining attention as alternatives to animal testing^46^. As such, closed-loop BNN platforms could serve as versatile platforms for modeling and investigating the responses of living neurons under complex, task-driven conditions, extending beyond spontaneous activity paradigms. Finally, BNNs could also serve as novel computational resources, replacing the current power-demanding hardware that is used for machine learning^47^. While many challenges remain (see the Discussion section in Ref.^12^), BNNs could offer multiple advantages, such as adaptability^12,21^, scalability, energy efficiency, and the opening of possibilities for developing “wetware” computing devices^48^.

## Materials and Methods

### Neuronal Culture

Primary cortical neurons were isolated from the cerebral cortices of embryonic day-18 (E18) rats. The cerebral cortices dissected from the rat embryos were placed in a 60-mm dish containing 4.5 mL of Hanks-balanced salt solution (HBSS; Gibco 14175-095) supplemented with 10 mM HEPES (Gibco 15030-015) and 1% penicillin/streptomycin (Sigma P-4333). The tissues were minced into approximately 1 mm^3^ pieces and transferred to a 15-mL centrifuge tube together with the HBSS. To the tube, mL of 2.5% trypsin (Gibco 15090-046; final concentration: 0.25%) and 0.2 mL of 10 mg/mL DNase (Roche 10104159001; final concentration: 0.4 mg/mL) were added. The mixture was then incubated at 37 °C for 15 minutes.

Following the incubation step, the solution was completely aspirated, and the tissue was rinsed three times with a fresh HBSS. The tissue was then mechanically dissociated by trituration with fire-polished glass pipettes to obtain a single-cell suspension.

Before cell plating, the electrode surface was coated with polyethyleneimine (PEI) and laminin to promote cell adhesion. First, a 1% terg-a-zyme (Alconox 1304-1) was applied to the surface and left at room temperature for 2 h. After this, the chip was thoroughly rinsed three times with deionized water. The chip was then submerged in a beaker filled with 70% ethanol and left at room temperature for 30 min. Subsequently, the chip was thoroughly rinsed with deionized water and dried. A 0.07% PEI/borate buffer solution was then applied to cover the entire electrode surface and incubated for 1 h. Next, the surface was rinsed three times with deionized water, dried, and coated with a 50–100 µg/mL laminin/DPBS solution in the same manner as that used for the PEI solution. Finally, the surface was rinsed three times with deionized water, dried, and prepared for further steps.

After the coating, the microfluidic device was attached to the HD-MEA chip by placing it over the electrode area with forceps. After allowing it to attach overnight, the chip well was filled with neuronal plating medium consisting of minimum essential medium (Gibco 11095–080) supplemented with 5% fetal bovine serum (Gibco 12483) and 0.55% glucose (Sigma G-8769). The chip was then degassed in a vacuum chamber to remove the air bubbles entrapped in the microfluidic device. The medium was then exchanged with fresh neuronal plating medium, and the chip was incubated in a CO_2_ incubator (37 °C, 5% CO_2_) for at least one night before cell seeding.

Cortical neurons were seeded onto the HD-MEA chip at a density of 700 cells/mm^2^. One hour after plating, the medium was completely replaced with Neurobasal medium (Gibco 21103–049) supplemented with 2% B-27 (Gibco 17504–044) and 1% GlutaMAX-I (Gibco 35050–061). Half of the medium was subsequently replaced with fresh Neurobasal medium twice per week. Recordings were performed using neurons at 14–31 DIV.

The neuronal growth observed on the HD-MEA chips was assessed by labeling the cells with the fluorescent chemical probe NeuO^49^. First, half of the medium (500 µL) was removed from the well and mixed with 1.5 mL of fresh Neurobasal medium. The resulting mixture was split into two 1-mL aliquots. To one of the tubes, 2 µL of 100 µM NeuO solution was added, and the mixture was thoroughly stirred (STEMCELL Technologies 01801; final concentration: 2 µM). Subsequently, the remaining medium in the well was completely aspirated and replaced with NeuO-containing medium. After a 30-min incubation period, the medium was completely replaced with NeuO-free medium, which was incubated at 37 °C. Observations were acquired via a stereomicroscope (Nikon SMZ18) equipped with an sCMOS camera (Andor Zyla 4.2P).

### Microfluidic Devices

The microfluidic devices for patterning primary neurons were fabricated as described previously^20,39^. Briefly, a master mold was fabricated by patterning an SU-8 photoresist on a silicon wafer, and the PDMS gel was poured onto the mold and thermally cured in a 70 °C oven for 2 h. The hardened PDMS microfluidic device was then peeled off the mold and cut with a clean razor blade. The device was then cleaned via sonication in ethanol for 5 min, rinsed in ddH_2_O water three times, sterilized under UV light for 30 min, and finally attached to the recording area of the MaxOne HD-MEA chip as described in the previous section.

Each device consisted of 128 square wells, each with a side size of 100 µm, which were separated by 100 µm. The wells were connected with microtunnels in either a lattice or modular configuration (see Fig. 1C). The microchannels were fabricated in two configurations that varied in height: one with a width of 5.1*±*0.5 µm and a height of 5.4*±*0.2 µm (mean*±*SD, *n* = 28 channels) and another with a width of 5.4*±*0.7 µm and a height of 1.2*±*0.04 µm (*n* = 8 channels). Both devices were employed in experiments without performing systematic selection. The overall size of the microfluidic film, including the boundary regions, was approximately 3.3 mm by 1.7 mm.

### HD-MEA Recording

Neuronal recording and stimulation were performed via the MaxOne HD-MEA system (MaxWell Biosystems). The MaxOne+ chip is a device featuring 26,400 Pt electrodes (without a Pt-black coating) arranged at a 17.5 µm pitch within a recording area of 3.85*×*2.10 mm^2^. The extracellular potential was recorded, and its high sampling rate of 20 kHz enabled the production of an excellent temporal resolution. Signals with amplitudes exceeding five times the standard deviation of the amplitude of the preceding signal were detected as spikes. Furthermore, electrical pulse stimulation could be delivered via a subset of electrodes, with a maximum of 32 electrodes. All processes related to the recording and stimulation were implemented via Python-based custom scripts.

To identify “active electrodes” that were capable of capturing reliable neural activity, the spontaneous activity level was measured for 30 s across all electrodes within the square-well region. Thresholds were then applied to the mean firing rate and mean firing amplitude of each electrode to select the active electrodes (less than 1,024), which were used for recording purposes. From the selected recording electrodes, the top 10% of electrodes (on the basis of their mean spike amplitudes) were identified. Of these, 30 electrodes were randomly selected to serve as the electrodes for delivering feedback stimulation during closed-loop control. These stimulation electrodes were further divided into two groups based on the polarity of the output values produced for the periodic signal learning task.

### Closed-Loop Control

A closed-loop system was realized by controlling the HD-MEA system through a Python/C++ API. The entire closed-loop control scheme operated on a cycle with a mean duration of 332.5*±*1.5 ms (mean*±*SD; *n* = 138 recordings from 46 cultures) and an SD of 3.7*±*0.7 ms. Each cycle consisted of a 120 ms pause before starting spike counting to process the waveform through an FIR filter and remove stimulation artifacts, a spike accumulation period of 200 ms, and software/hardware latencies.

Two tasks were considered in this study, i.e., a sine wave generation task and a Lorenz attractor generation task. For the former task, the target signal *d* was set to

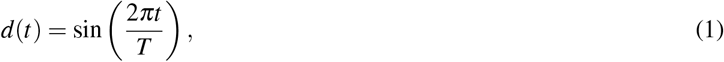

where *T* (= 4, 10, and 30 s) is the cycle period. For the latter task, we employed the classic Lorenz attractor given by

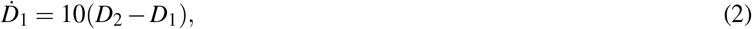

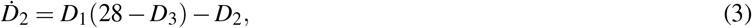

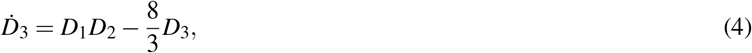

and its affine transformation 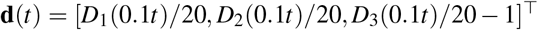was then used as the target signal. The system exhibited deterministic chaos, and the phase space revealed the strange attractor illustrated in Fig. 5A. The value of the target waveform was sampled at each cycle; thus, the experiment was susceptible to fluctuations in the intervals of feedback cycles.

A schematic representation of the system is summarized in Fig. 1A. The system consisted of four modules, including the HD-MEA, each with the following functions:

- **Spike Counting (C++)**: Received spike information concerning the specified electrodes from the hardware in real time.
- **Data Routing (Python)**: Accumulated the spike counts of each electrode over a specified duration (set to 200 ms) and transferred the data to the subsequent modules.
- **Reservoir Computing (C++)**: Filtered the spike trains with a double exponential filter to generate the reservoir state vector **x** = [*x*_*i*_]^36^:

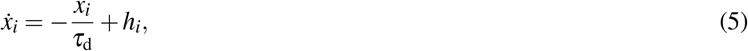

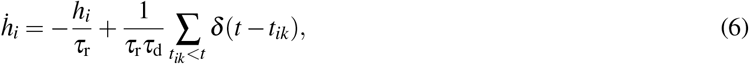

where *i* is an index for neurons (electrodes), *h* is an auxiliary variable, and *τ*_r_(= 500 ms) and *τ*_d_(= 2000 ms) are rise and decay time constants, respectively. *t*_*ik*_ is the *k*th spike fired by the *i*th neuron, and *δ* is the Dirac delta function with *δ* (0) = 1. The module then computed the output vector as follows:

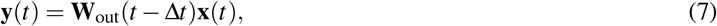

and optimized the output weight matrix **W**_out_ for use in the next time step. **W**_out_ was initialized as zero and optimized online via the FORCE learning algorithm^4^:

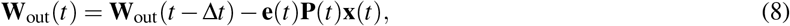

where **e** and **P** are the error vector and the inverse correlation matrix, respectively, which are calculated as follows:

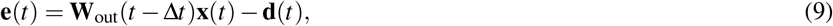

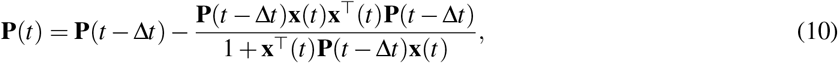

where **d** is the target signal vector. The initial value for **P** was set to

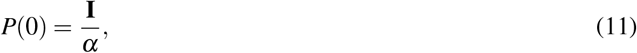

where *α* is the “learning rate” set to 1000 and **I** is an identity matrix.
- **Stimulation Sequencer (Python)**: Produced commands for delivering feedback stimulation on the basis of the output and sent them back to the MEA hardware (MaxOne Hub). Biphasic voltage pulses, each with a duration of 200 µs per phase, were applied with amplitudes determined according to the reservoir output **y**(*t*). Stimulation electrodes were selected on the basis of the procedure described in the ‘HD-MEA Recording’ section.

For the sine wave generation task, the amplitude of the applied biphasic pulse *A*(*t*) was determined as follows:

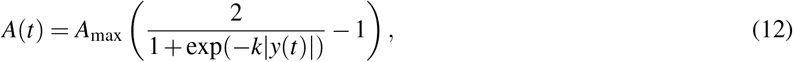

where *A*_max_ (= 876 mV) is the maximum amplitude, *y* is the reservoir output, and *k* (= 4) is the gain for adjusting the nonlinear relationship between *y* and *A* (Fig. 3A). The parameters were set on the basis of pilot experiments examining the relationship between the pulse amplitude and evoked activity (Fig. 3B). The stimulation electrodes were divided into two groups. For the first group, the stimulation amplitude was set to *A*(*t*) when *y*(*t*) *≥* 0 and to 0 when *y*(*t*) *<* 0. For the second group, the amplitude was set to *A*(*t*) when *y*(*t*) *≤*0 and to 0 when *y*(*t*) *>* 0.

For the Lorenz attractor generation task, the stimulation electrodes were divided into three groups, each corresponding to one of the three components of the signal. The output vector **y** = (*y*_1_, *y*_2_, *y*_3_) was then mapped onto the amplitude of the feedback stimulation as follows:

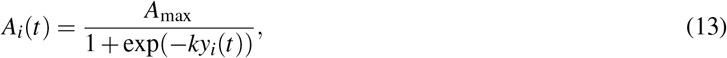

where *i* is the index for the signaling component, *A*_max_ (= 876 mV) is the maximum amplitude, and *k* (= 4) is the gain.

## Data Analysis

### Spontaneous Activity

The extracellular potentials recorded from each electrode were filtered (1–3000 Hz) and recorded in an HDF5 file, together with the spikes that were detected in real time by thresholding the extracellular signals at five times the standard deviation. The spikes were then binned at 1 ms and filtered with the same double exponential filter described above to obtain a (virtual) firing rate, or a “reservoir state” in the case of the closed-loop experiments (see the next section).

The Pearson correlation coefficients *r*_*ij*_ between electrodes *i* and *j* were subsequently computed via the following equation:

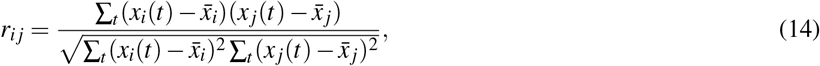

where *x*_*i*_(*t*) and *x* _*j*_(*t*) are the firing rates of electrodes *i* and *j* at time *t*, respectively, and 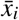 and 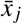 denote the mean firing rates of electrodes *i* and *j*, respectively.

PCA was used to extract the underlying, low-dimensional dynamics that govern the activity of a large population of neurons. For a recording with a firing rate matrix of **X***∈* ℝ^*T×d*^, where *T* and *d* are the numbers of time points and electrode channels, respectively, **X** was first centered on its mean as follows:

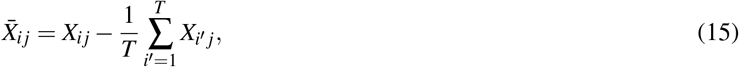

with *X*_*ij*_ denoting the elements of **X**. Denoting the centered firing rate matrix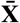, factor loadings and the variance explained by the PCs were then obtained as an eigendecomposition of the covariance matrix, 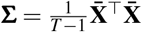:

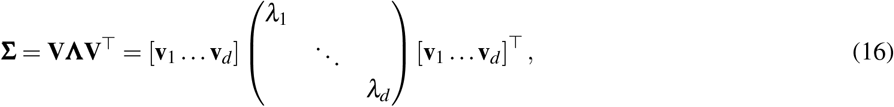

where **V** is the eigenvector matrix and **Λ**is the diagonal matrix of the eigenvalues *λ*_*i*_. The eigenvectors **v**_*i*_ correspond to the factor loadings of the *i*-th PC, and each normalized eigenvalue 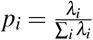 is the variance explained by the corresponding PC. To quantify the effective dimensionality of the network dynamics, the effective rank of **X** was calculated as follows^37^:

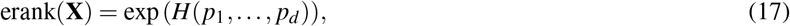

where *H* is the Shannon entropy given by

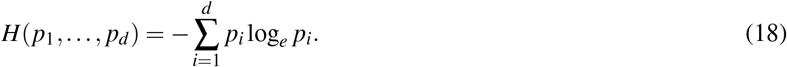

### Closed-Loop Experiments

The network dynamics produced under closed-loop control were analyzed following the same procedure used for spontaneous activity, except that the stimulation artifacts needed to be removed. Specifically, the artifacts were removed by excluding all spikes occurring within 10 ms before and after the stimulation.

To characterize the trajectory of the neural dynamics, the activity observed under closed-loop control was projected into a PC space derived from the spontaneous activity. Let 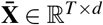 denote the centered reservoir state matrix during a closed-loop experiment, and **V**^*\**^ = [**v**_1_ … **v**_*k*_] denote a submatrix of the eigenvector matrix derived from the spontaneous activity. The neural activity trajectory projected onto a *k*-dimensional PC space, i.e., **X**^*\**^ *∈* R^*T×k*^, was then calculated as follows:

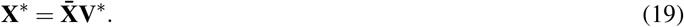

The analyses were performed in MATLAB (MathWorks).

## Acknowledgments

The authors thank Prof. Wilten Nicola at the University of Calgary for discussion on the inherent properties of the FORCE learning algorithm. We also thank Mr. Iori Morita at Tohoku University for the fabrication of the photomasks. This work was partly supported by MEXT Grant-in-Aid for Transformative Research Areas (A) “Multicellular Neurobiocomputing” (24H02330, 24H02332, 24H02334), JSPS KAKENHI (22H03657, 22K19821, 22KK0177, 23H00251, 23H02805, 23H03489, 25H00447), JST CREST (JPMJCR19K3), JST ALCA-Next (JPMJAN23F3), the WISE Program for AI Electronics of Tohoku University, and the Cooperative Research Project Program of the RIEC, Tohoku University. This research was carried out at the Laboratory for Nanoelectronics and Spintronics, RIEC, Tohoku University.

## Supplementary Figures

**Fig. S1.**
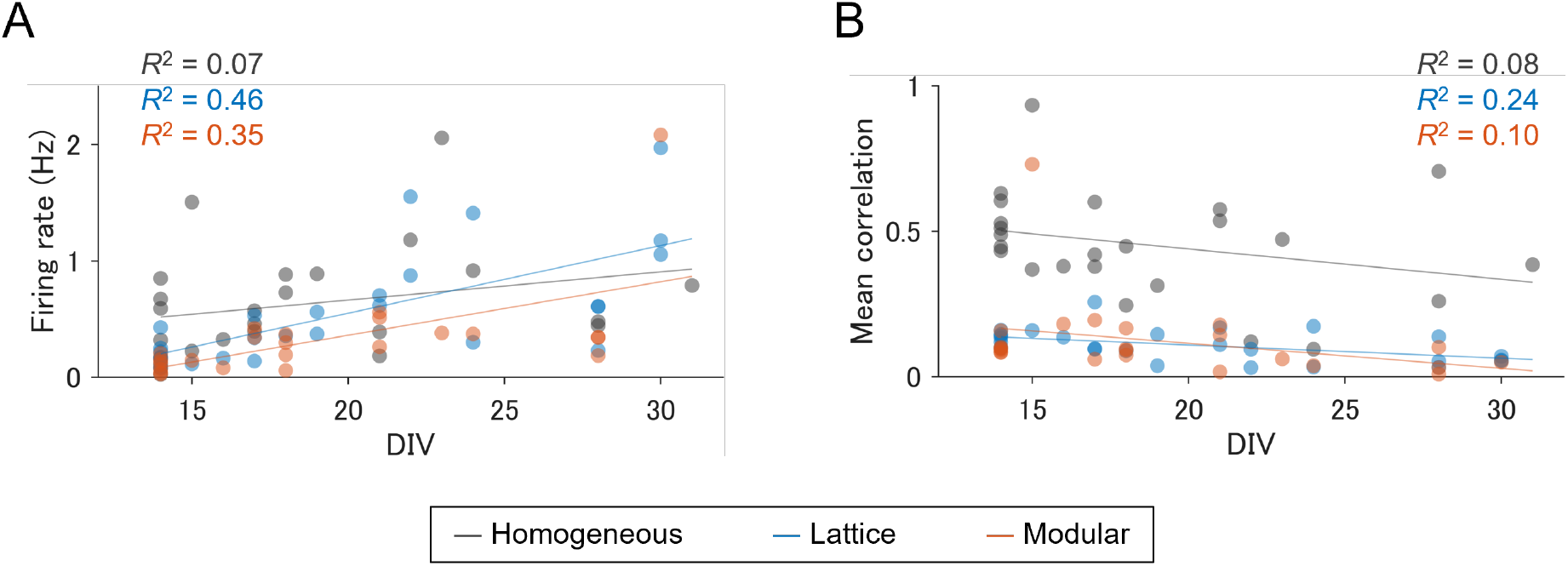
Dependence of network statistics on culture age. (A) Mean population firing rate and (B) mean pairwise correlation are plotted as a function of days in vitro. While lattice and modular networks (blue and orange, respectively; *n* = 27, 24) exhibited a stronger dependence on culture age compared to homogeneous networks (gray, *n* = 24), the overall dependence remained weak, with the coefficient of determination (*R*^2^) below 0.5 in all cases.

**Fig. S2.**
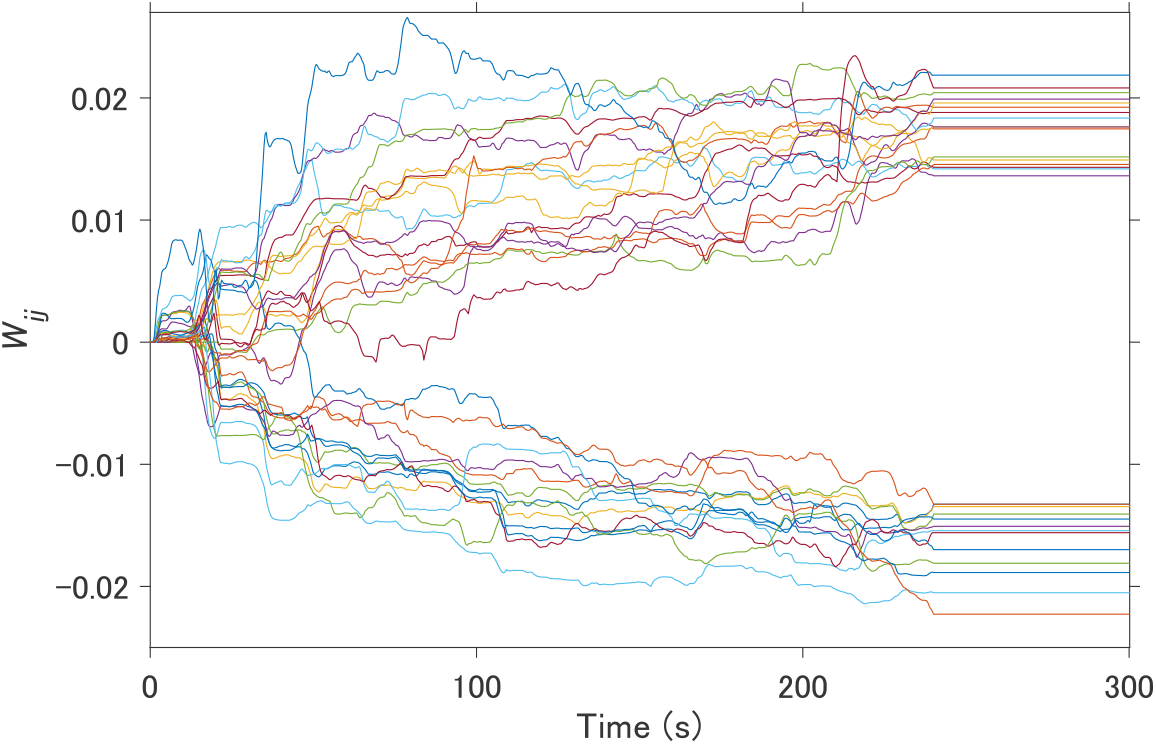
Evolution of readout weights during training. Readout weights were trained over a period of 0 to 240 s. The top 20 elements with the highest final values were selected, and their temporal profiles plotted. Data shown are for a modular network at DIV 23, from the same experimental run presented in Fig. 3A of the main text.

**Fig. S3.**
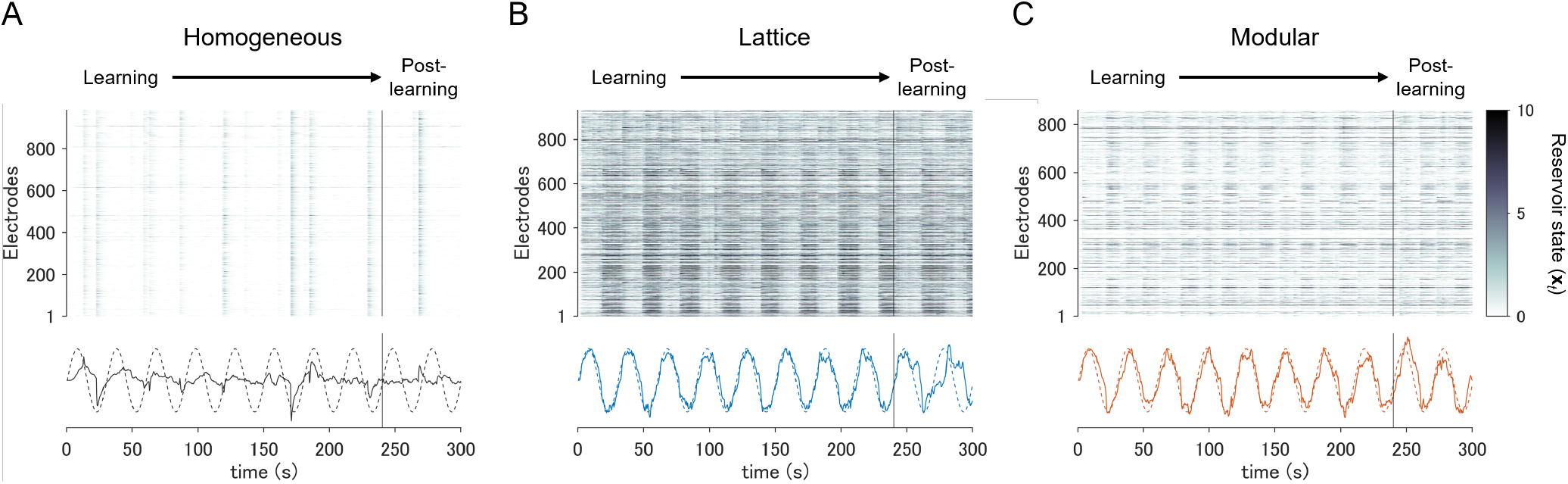
Effect of network morphology on task performance. Filtered raster plots (reservoir states, **x**_*t*_) and corresponding output signals for (A) a homogeneous network at DIV 21, (B) a lattice network at DIV 24, and (C) a modular network at DIV 23. The target signal (dashed curve) was a sinusoidal wave with a period of 30 s. Fig. 4C of the main text is reproduced here as panel B for clarity.

